# Combined SPT and FCS methods reveal a mechanism of RNAP II oversampling in cell nuclei

**DOI:** 10.1101/2022.07.27.501703

**Authors:** Marie Fournier, Pierre Leclerc, Aymeric Leray, Dorian Champelovier, Florence Agbazahou, Gabriel Bidaux, Alessandro Furlan, Laurent Héliot

## Abstract

Gene expression orchestration is a key question in fundamental and applied research. Different models for transcription regulation were proposed, yet the dynamic regulation of RNA polymerase II (RNAP II) activity remains a matter of debate. To improve our knowledge of this topic, we investigated RNAP II motility in eukaryotic cells by combining Single Particle Tracking (SPT) and Fluorescence Correlation Spectroscopy (FCS) techniques, to take advantage of their different sensitivities in order to analyze together slow and fast molecular movements. Thanks to calibrated samples, we developed a benchmark for quantitative analysis of molecular dynamics, to eliminate the main potential instrumental biases. We applied this workflow to study the diffusion of RPB1, the catalytic subunit of RNAP II. By a cross-analysis of FCS and SPT, we could highlight different RPB1 motility states and identifyed a stationary state, a slow diffusion state, and two different modes of subdiffusion. Interestingly, our analysis also unveiled the oversampling by RPB1 of nuclear subdomains. Based on these data, we propose a novel model of spatio-temporal transcription regulation. Altogether, our results highlight the importance of combining microscopy approaches at different time scales to get a full insight into the real complexity of molecular diffusion kinetics in cells.

## INTRODUCTION

Biological activities are classically modulated through the dynamic binding and unbinding of molecular partners ^1^. Depending on the process considered, molecules may exist in several distinct complexes, which differ in their motility and availability features. In the frame of gene expression modulation, diffusion and target search constitute critical features for transcription actors ^2,3^. It was acknowledged for a long time that RNA Pol II was recruited on the target genes by general transcription factors during pre-initiation complex (PIC) formation ^4–6^. Indeed, RNAP II was initially observed in mammalian cells in cluster-like structures by immunostaining of the polymerase active form in fixed cells and was co-localized with nascent RNAs ^7,8^. This led to the interpretation of static transcription factories. Recent advancements in the field however suggest that RNA Pol II assembly is very dynamic throughout time and that several RNA pol II can be co-recruited and form transient clusters which disintegrate after a few seconds, depending on the transcriptional activity ^9–11^. How these clusters match with the concepts of transcription factories or TAD (Topologically Associated Domains)^12^, and how they organize over time remain elusive. In light of these results, it is interesting to revisit the investigation of RNAPII mobility by taking advantage of the progress in molecular labelling as well as in camera speed and detector sensitivity.

Various microscopy approaches have been developed in the last decades to quantify and/or track molecular motilities. Fluorescence microscopy has notably been widely used to decipher the dynamics of molecular orchestration and is a key element to push forward the understanding of spatiotemporal molecular mechanisms. The mobility of a given molecule in cells classically encompasses several diffusion modes ^13,14^, and retrieving quantitative data about the diffusion of both fast and slow molecules is critical to understanding the dynamics of molecular events, including transcription, in the nucleus. To this end, we took advantage of the complementarity of FCS (Fluorescence Correlation Spectroscopy) high temporal resolution and SPT (Single-Particle Tracking) accuracy in spatial measurements of slower displacements, to perform a more complete analysis of RNAP II dynamics.

The calculation of molecular scattering using FCS is based on the autocorrelation of the measurement of the fluorescence fluctuations due to the transit of fluorescent molecules through the excitation focal volume of a laser beam ^15^. Although the focal volume is small (about 0.1 μm^3^), it corresponds at the nucleus scale to a subdomain that can encompass several distinct molecular crowding conditions or molecular interactions, and each experimental point thus already constitutes an average of several behaviors ^16^. Moreover, such fluctuation-based techniques are blind to immobile molecules ^17,18^, which constitutes a drawback that has to be taken into account. FCS is rather suitable for processes occurring faster than 1 μm^2^/s ^21^.

In parallel, super-resolution SPT techniques have been developed thanks to the improvement of fluorophores and camera sensitivity over the last decade, and allow tracking of individual molecules in the field of view over time ^19,20^. Their temporal sampling rate (classically 10 ms/frame for a 10 μm^2^ area) allows the measurement of molecular motilities with diffusion coefficients up to 5 μm^2^/s ^14,21^. In addition, SPT techniques require the analysis of a large number of individual traces (10^3^-10^5^ trajectories) that may not be equally represented as a function of their length, depending on the mobility of the molecules and the experimental set-up. These biases may result in an over-representation of slower molecules with longer trajectories and in less accurate statistics concerning faster molecules, which is often underestimated in SPT analyses ^3^.

In this study, we first evaluated the results obtained by FCS and SPT methods with reference samples, to establish a benchmark for the use of these methods to unravel RPB1 behavior in the nucleus. This allowed us to overcome the potential biases that could derive from the different timescales and different analytical protocols of the two techniques. We reasoned that it was important to deal with experimental data rather than simulations because the former describe at best the reality of experimental images and instrumental constraints and contributions. We came up with a robust process displaying optimized conditions, which we applied to the analysis of RPB1 dynamics. Thanks to it, we show that RNAP II dynamics in living cells can be described by at least four different mobility modes with a significant proportion of anomalous diffusion probably corresponding to enhanced local target search by RNAP II.

## RESULTS

Before applying FCS and SPT techniques to investigate RNAP II dynamics, we first characterized their sensitivity in our set-ups, thanks to a system based on the diffusion of fluorescent beads in different media of controlled viscosity. To recapitulate the apparent diffusion values observed in the literature for proteins of the transcription complex in the nucleus, i.e. between 0.1 and 15 μm^2^/s ^3,22^, we selected appropriate parameters to perform FCS and SPT experiments (see Material and Methods for more details).

### Characterization of calibrated bead diffusion by ACF analysis of FCS measurements

The FCS experiments were conducted with fluorescent microspheres in a water/glycerol mix, with a proportion of glycerol ranging from 0 to 80 %. These combinations provided us with control solutions displaying theoretical diffusion coefficients ranging from 1 μm^2^/s to 15 μm^2^/s (Table S1).

We chose to fit the autocorrelation curves resulting from these experiments with an anomalous model, which is classically used when studying the molecular motility in living cells ^23–25^, and allowed us to maximize the dispersion range of the results and to explore at the same time the goodness of this model that we would use later to investigate RNAP II diffusion in cells. We globally obtained a good correlation between the theoretical values and our FCS experimental values (Fig.1 and Table S1).

**Figure 1.**
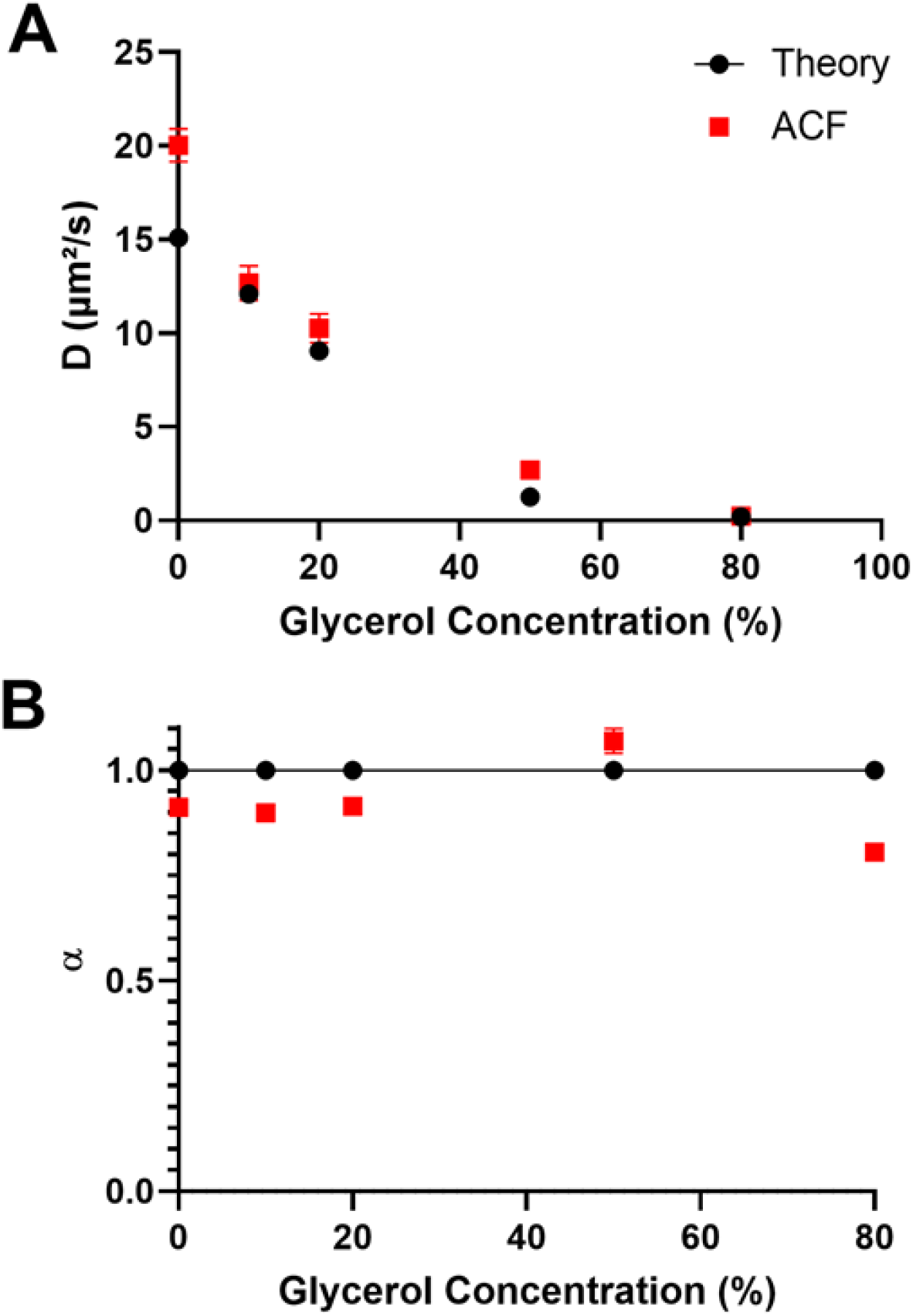
Characterization of diffusion by Fluorescence Correlation Spectroscopy. Fluorescent microspheres were resuspended in water/glycerol mixtures, with proportions of glycerol ranging from 0 to 80 %. The ACF curves obtained from FCS acquisitions were fit with an anomalous model to infer the anomalous coefficients (A) and diffusion coefficients (B). Experimental values (red squares) can be compared to theoretical values (black dots). Experimental results are expressed as means +/− SD from at least 20 independent measurements.

We noticed a slight overestimation of our computed results for the diffusion coefficients when compared to the theoretical values, especially for the diffusion of beads in pure water (Fig.1A). This could stem from a minimal error in the estimated value of viscosity inferred, as previously reported by ^26^ and described in Material and methods. Altogether, we could assess diffusion coefficients ranging from 1 to 15 μm^2^/s in our FCS set-up.

Besides, up to a glycerol content of 50 % in water, the ACF analysis yielded an anomalous coefficient value very close to the theoretical value of 1 (between 0.96 and 1.02), as expected for Brownian diffusion (Fig.1B). Only the diffusion of beads in the solution containing 80 % glycerol was described as anomalous, with an alpha of 0.70 +/− 0.02. However, it is noteworthy that this mixture might be non-perfectly homogenous due to the high viscosity of glycerol, and that may result in anomalous diffusion, as previously described in the literature ^27,28^.

### Characterization of calibrated bead diffusion by SPT analysis

To compare the measurements by FCS and SPT, we applied a similar strategy to analyze the diffusion of reference beads with SPT.

We performed SPT acquisitions in HILO (Highly Inclined and Laminated Optical sheet) microscopy mode ^29^, a technique classically used for nucleus imaging because it allows a selection of the plan of observation thanks to a very inclined optical sheet. We analyzed image series with the MTT (Multiple-Target Tracing) algorithm tracking software ^30^ to detect and reconstruct trajectories of the particles. Since short trajectories strongly penalize the stability of the adjustment function methods ^31^, we only selected trajectories with more than 10 detections, which are more reliable. With this cut-off, we obtained a mean number of detections per trajectory superior or equal to 20 for all conditions (Fig.S1). In concordance with FCS analysis, the trajectories were analyzed with an anomalous model. We obtained the diffusion parameters from the linear regression of the logarithmic MSD (Mean Square Displacement) of all trajectories as a function of time, alpha being the slope of the line and D_ɑ_ the value at the origin. We found a computed anomalous coefficient (ɑ) very close to 1, which is consistent with the fact that beads in solution should present a Brownian movement (Fig.2A). As observed in FCS, a high content of glycerol (80%) resulted in an alpha computed at 0.8, with this apparent anomaly probably associated with a certain heterogeneity of the mixture.

**Figure 2.**
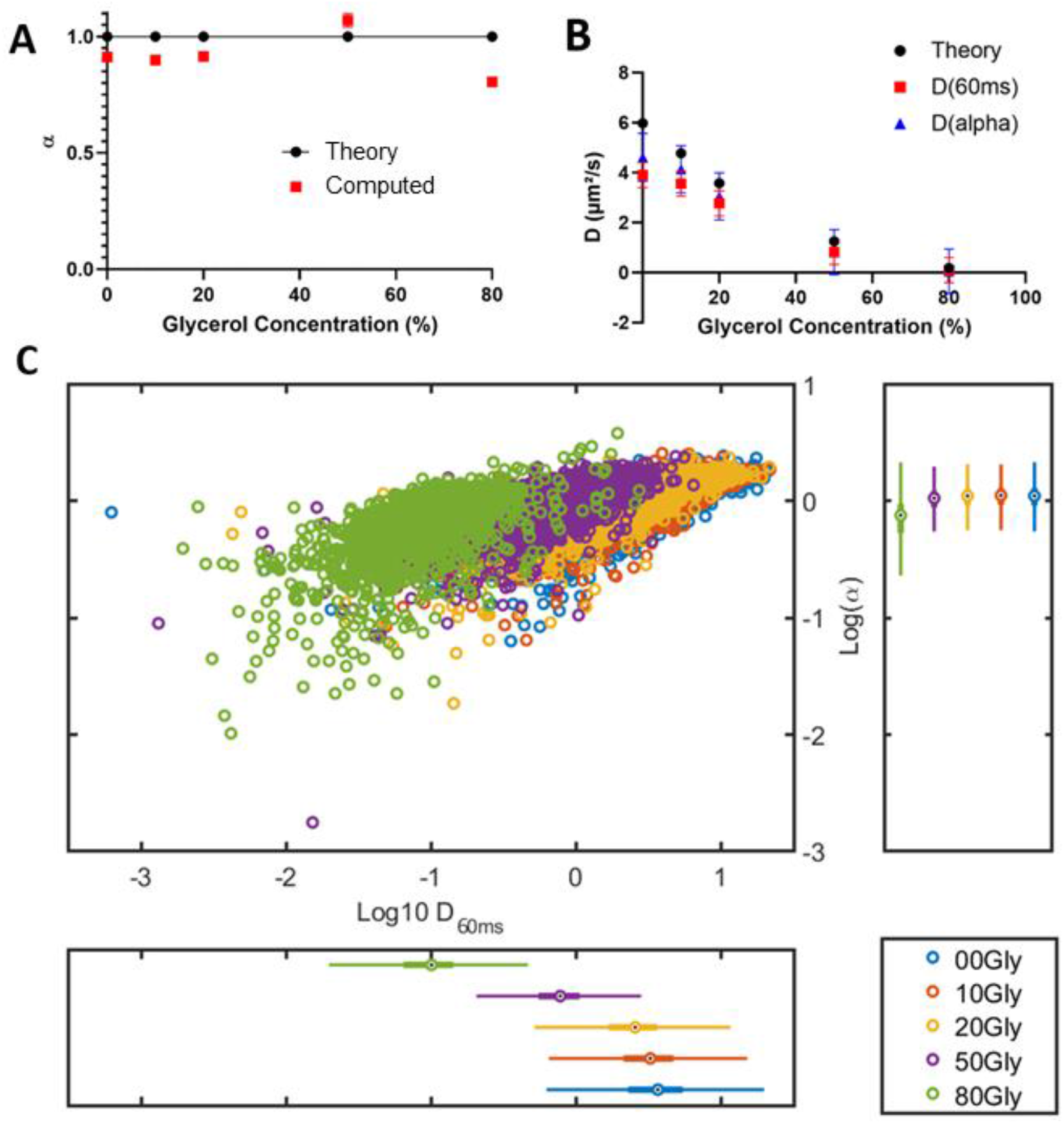
Characterization of diffusion by Single Particle Tracking. Fluorescent microspheres were resuspended in water/glycerol mixtures, with proportions of glycerol ranging from 0 to 80 %. SPT acquisitions were performed at a frame rate of 100 Hz. A. The anomalous coefficient α is provided as a function of the glycerol proportion. Experimental values (red squares) can be compared to theoretical values (black dots). B. The diffusion coefficients were computed by two methods: D_a_ (blue triangles) were obtained from the slopes of TEA-MSD (Time Ensemble Average Mean Square Displacement) expressed as log, while D_60ms_ were calculated for each trajectory during the first 60ms of its occurrence (red squares). These experimental values could be compared to theoretical values (black dots). Both methods yielded an asymptotic trend for high D values. C. The distributions of D_60ms_ and α from the different experimental conditions are represented with log scales in a scatter plot. Distinct subpopulations can be distinguished (green, purple, and yellow dots) within a limited range of parameters, whereas some others overlap (yellow, red, and blue dots). Data come from at least one hundred measurements per condition.

At the same time, we computed the diffusion coefficients D_ɑ_ from the linear regression of MSD, as explained in the previous paragraph. Besides, since it is assumed that, at short time intervals, molecules behave like Brownian particles, we also calculated the diffusion coefficient D_60ms_ for each trajectory by considering the slope of the vector from the origin to its value at 60 ms, as previously performed in the community ^32^.

Both methods yielded diffusion coefficients slightly smaller than the theoretical values (Fig.2B), as previously described and discussed ^33^. Interestingly, the standard deviation associated with D_60ms_ was two times lower when compared to that of D_ɑ_, so we chose the D_60ms_ method for our subsequent SPT analyses of RPB1 diffusion.

We noticed that, for theoretical values of diffusion coefficients above 4 μm^2^/s, the values of D_60ms_ and D_ɑ_ were lower than the expected theoretical values, suggesting that we had reached a plateau (Fig.2B). Thus, this calibration step shows that SPT allows to study molecular diffusion below 4 μm^2^/s. This agrees with the classical measurement ranges observed in the literature with this method, due to limitations in camera frame rates, tracking algorithms and fluorophore brightness.

Consistently, the representation of measurements as a scatter plot (Fig.2C) nicely illustrated the fact that the populations that we detected at different glycerol concentrations could be easily distinguished, up to this plateau where they overlapped. The distribution of bead diffusion measurements still evidenced that this method was well suited to detect particles diffusing with coefficients over several orders of magnitude.

To sum up this characterization step, in our experimental conditions, FCS was able to detect relatively high (in the biological field) diffusion coefficients, ranging from 1 to 15 μm^2^/s, while SPT allowed us to detect slower diffusions, ranging from 10^−2^ to 4 μm^2^/s, thus confirming the complementarity of these methods.

### Analysis of SPT data including mixed populations

In order to evaluate how our approach could discriminate mixed subpopulations in a sample, as is the case in cells, we constituted mixtures of bead trajectories within various glycerol contents, from the previous data. We chose to use the histogram analysis of MSD (called h-MSD), which allows to get a comprehensive overview of diffusion features in complex samples with a histogram representation based on the MSD analysis of each trajectory ^32^.

Under the assumption that D_60ms_ and (ɑ) distributions follow sums of power laws, one can discriminate different subpopulations emerging from a heterogeneous sample, by using a Gaussian Mixture Model (GMM) to fit the data of log(D) and log(ɑ) distributions. Using a methodology derived from unsupervised machine learning, we used the BIC (Bayesian Information Criteria) and AIC (Akaike Information Criteria) to infer the proper *k* number of populations (ranging from 1 to 10) by analyzing the point of inflection of the BIC and AIC curves as a function of *k*. Then we computed GMM with this *k* to fit our distributions.

We tested this methodology by mixing batches of data originating from distinct bead acquisitions with various glycerol contents and assessed its ability to discriminate subpopulations and retrieve both their right proportions and accurate coefficient values. Consistently, the BIC and AIC analyses indicated that two-population models best described our different mixtures (Fig.3A and B for a mixture of trajectories from the 0 % and 50 % glycerol conditions). We showed that the h-MSD/GMM method allowed for discriminating distinct subpopulations, with good quality in retrieving both the proper values and proportions for each subpopulation (Fig.S2 and Table S2). Logically, in these settings in which particles correspond to Brownian particles, the discrimination happened in the diffusion coefficient dimension while alpha logically remained centered around 1 (Fig.3C). This method was pushed towards its limits only when mixing subpopulations that were very close to the upper limit of detection, namely when merging populations with diffusion coefficients centered around 3.8 and 4.0 μm^2^/s, that were considered as a single population (Table S2D).

**Figure 3.**
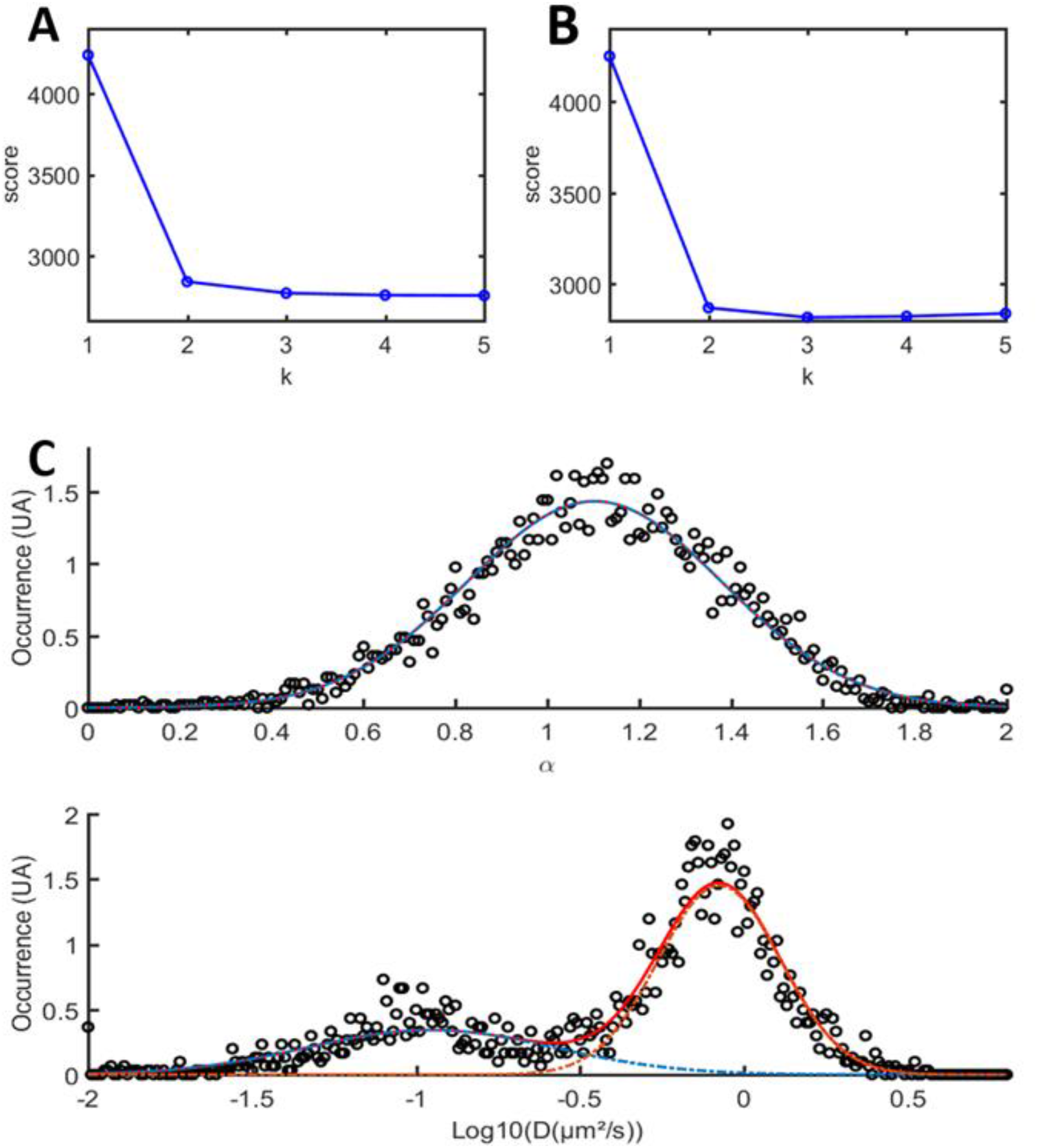
Discrimination of merged subpopulations thanks to h-MSD. SPT trajectories from experiments with fluorescent microspheres diffusing in 20 % and 50 % glycerol solutions respectively, were merged and submitted to the h-MSD analysis workflow to compute for each trajectory the alpha and log(D). The number of Gaussian functions to be found in our distribution was inferred from the AIC (Akaike Information Criteria) (panel A) and the BIC (Bayesian Information Criteria) (panel B) respectively. In this example, the inflection point occurs for a number of populations k=2, consistently with the mixture of two experiments. Using the results of the h-MSD analysis, we represented in panel C the distributions of alpha (upper graph) and D (lower graph). The distribution was then fitted with a GMM (Gaussian Mixture Model). The details of all combinations and the comparison of calculated values with the initial values are provided in the Supplemental Material.

### FCS detects RPB1 anomalous diffusion in living cell nuclei

Once this benchmarking of molecular dynamics analyses was validated, we applied our methods to dissect the orchestration of transcription in eukaryotic cells. For that purpose, we engineered U2OS cell lines to replace endogenous RPB1, the catalytic subunit of RNAPII, with an SYFP2 fluorescently-labeled amanitin-resistant RPB1 version (Fig.4A), following the strategy previously used by Darzacq and collaborators ^9,34^.

**Figure 4.**
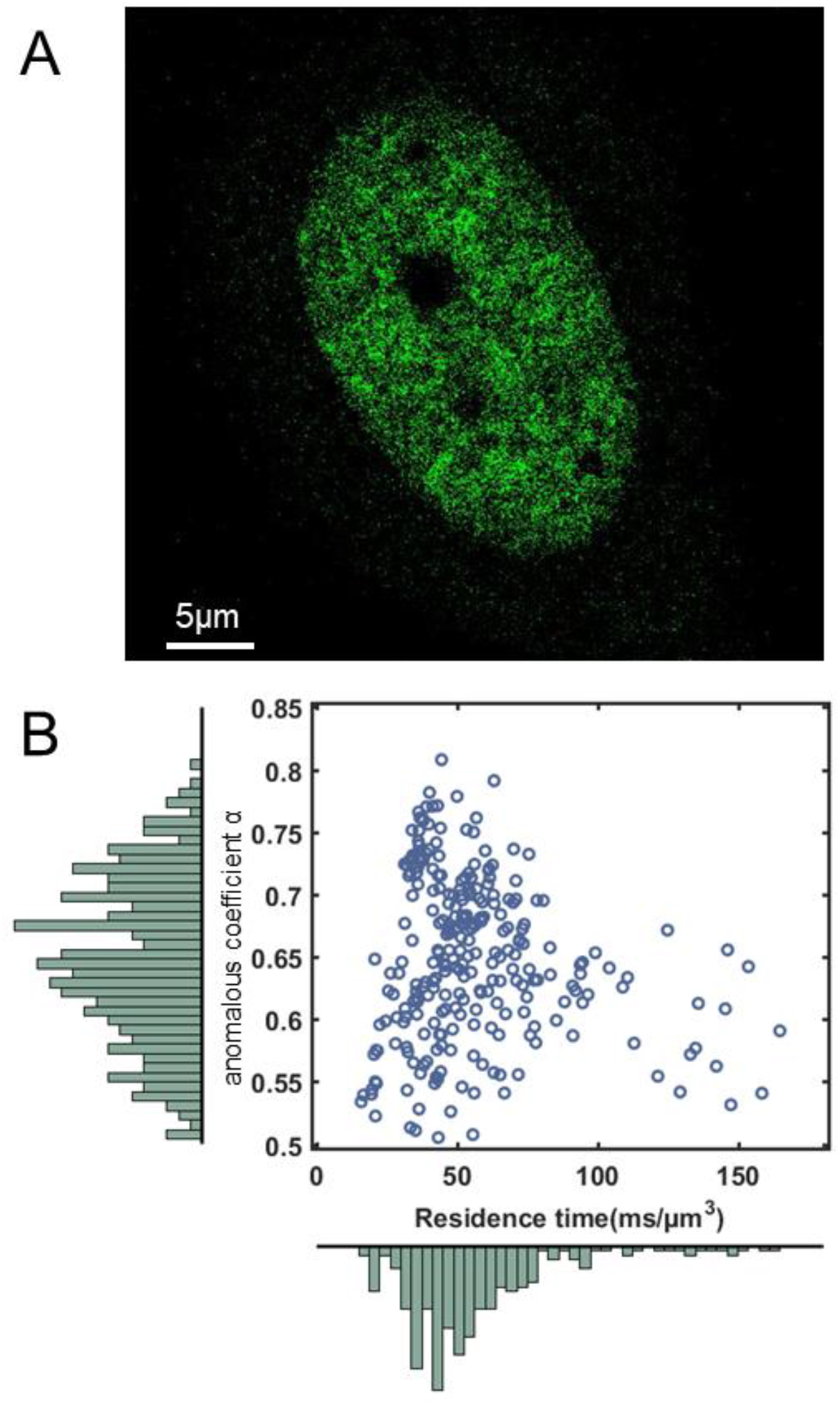
FCS detects RPB1 anomalous diffusion in living cell nuclei. A. Representative picture of a U20S RPB1-SYFP2 cell. B. Scatter plot of normalized residence times versus anomalous coefficients computed from 138 FCS measurements performed in U2OS cells expressing RBP1-SYFP2 (n=6). 80% of the transit times range from 23.7 and 89.1 ms/μm^3^ with an average of 61.9 ms/μm^3^ (D=5.7 μm^2^/s^α^). And the anomalous coefficients α range from 0.5 to 0.8 with a mean value of 0.61.

This allowed us to probe RPB1 diffusion by FCS. Anomalous coefficients were mostly distributed between 0.5 and 0.8, with a mean value of 0.61, indicating a restricted diffusion of RPB1 in nuclei (Fig.4B). Besides, we chose to compute the transit-times in the field of view (Tt, in ms/μm^3^) as a common metric to further compare the results between FCS and SPT (see material and methods for more details). Indeed, as described in the literature, it is not possible to directly compare the diffusion coefficients associated with different anomalous coefficients.

Most RPB1 transit times in FCS ranged from 20 to 150 ms/μm^3^, with a mean value of 62 ms/μm^3^ (D=5.7 μm^2^/s^α^). G(0) analysis determined that the number of RPB1 per focal volume is 2.6 +/−1.4 molecules.These transit times, as well as anomalous coefficients, were spread out as evidenced in the scatter plot, which is representative of a variety of different behaviors that could be detected by FCS. In addition, the distribution of these diffusion values was imperfectly fitted by a Gaussian model, suggesting the existence of subpopulations and/or the presence of some outliers.

### H-MSD SPT highlights the existence of distinct RPB1 subpopulations

We then performed SPT experiments to complete our analysis of RPB1 diffusion. SPT requires a dedicated labeling strategy to get only a limited number of molecules that can be tracked over time in the field of view. These molecules should be bright enough to be detected in a single particle tracking regime and possess a high photostability to be followed over a sufficient period (at least a few seconds, and ideally several tens of seconds). For that purpose, we took advantage of the novel Janelia fluorophores coupled to the Halo ligand JF549 ^35^ and engineered a stable cell line expressing RPB1 fused to the Halo tag.

In that context, we obtained staining levels adequate to identify single RPB1 molecules (Fig.5A), on which we were able to perform measurements of more than one thousand RPB1 trajectories (Movie 1 as an example) that were processed as aforementioned. We observed a great variety of molecular behaviors, as illustrated by the different kinds of tracks reported (Fig.5B), which shows the existence of various and complex behavior of RNAPII in the nucleus.

**Figure 5.**
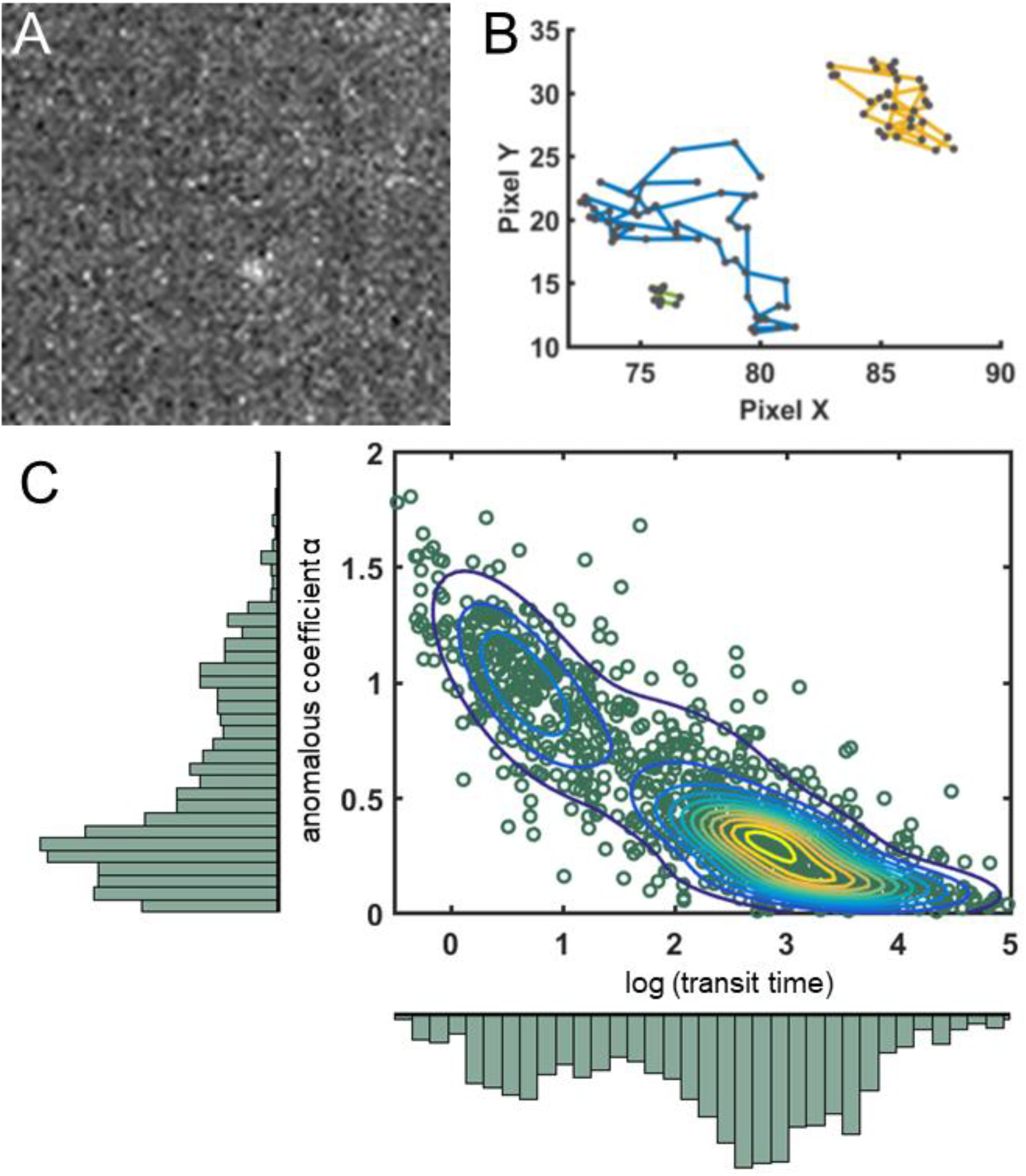
H-MSD SPT highlights the existence of distinct RPB1 subpopulations. A. Picture representative of the signal-to-noise ratio obtained with JF549-halo tag at 0.1nM in U2OS RPB1-Halo cells. B. Examples of trajectory obtained with the tracking pipeline. C. 1051 trajectories from SPT experiments carried out in the U2OS RPB1-Halo cell line (n=10), stained with JF549 at 0.1nM, were analyzed by h-MSD. Tt and α values from these experiments are represented in a scatter plot, in which concentric lines show the Probability Density Function given by the Gaussian Mixture Model. At least two distinct populations are clearly visible.

Following this workflow, we generated a scatter plot of anomalous coefficients and transit times corresponding to RNAP II (RPB1-Halo/JF549) trajectories (Fig.5C). The BIC and AIC analyses of the whole data suggested the presence of four distinct subpopulations (Fig.S3), but we limited our description to two main populations highlighted in the plot (Fig.5C). A majority one, probably encompassing several subpopulations as suggested by the distorted shape of the Probability Density Function, represented 75% of the trajectories, and was centered around a transit time Tt=777 s/μm^3^ (D=0.02 μm^2^/s) and an anomaly coefficient α=0.31. A second one, corresponding to 25 % of the trajectories, displayed a mean Tt=4.2 s/μm^3^ (D=0.5 μm^2^/s) and ɑ=0.95. In addition, around 20 % of RNAP II observed displayed a D_60ms_ < 5 10^−3^ ms/μm^2^ and were considered as immobile (Movie 2).

When comparing with FCS measurements (Fig.4B), one can observe that SPT allowed us to detect distinct diffusion behaviors since the smallest transit times detected in SPT exceeded the highest values measured by FCS (see also Fig.S4).

### Spatio-temporal localization of RPB1 subpopulations

Another intriguing observation that we made resides in the fact that no static regions with increased RPB1 concentrations could be observed within our super-resolution experiments. To get enough detections to infer the spatial information, we chose to stain the cells with a JF549 concentration higher than that used for multiple particle tracking and trajectory analysis. This level of signal allowed us to image the whole nucleus and investigate the global spatio-temporal distribution of RPB1. The analysis of detections along the acquisition indicated that spots appeared and disappeared over time, demonstrating the highly dynamic nature of these elements (Movies 3, 4 and 5).

When checking the mean intensity projection signal along the movie, we could observe that some areas were highlighted, that could correspond to transcription areas, over a diffuse background, showing that RPB1 explores the whole nucleus (except for nucleoli (Fig.6A). In this condition, tracking could be performed on several particles at the same time (Fig.6B). Moreover, when summing all the detections occurring throughout the acquisition, we noticed that some regions were more frequently visited by RPB1 molecules (Fig.6C), shedding light on a previously undescribed mode of dynamic spatio-temporal regulation of gene transcription by RNAP II oversampling. Importantly, when averaging the detections from all frames, we obtained a quite homogenous noisy background (data not shown), indicating that the regions highlighted in Fig.6A and B did not result from a static phenomenon. When comparing the signals from Fig.6A and 6C (merged in Fig.6D), one can see that regions with higher intensity mainly match, still, a great proportion of the nucleoplasm was visited by RPB1 but did not lead to detections by SPT.

**Figure 6.**
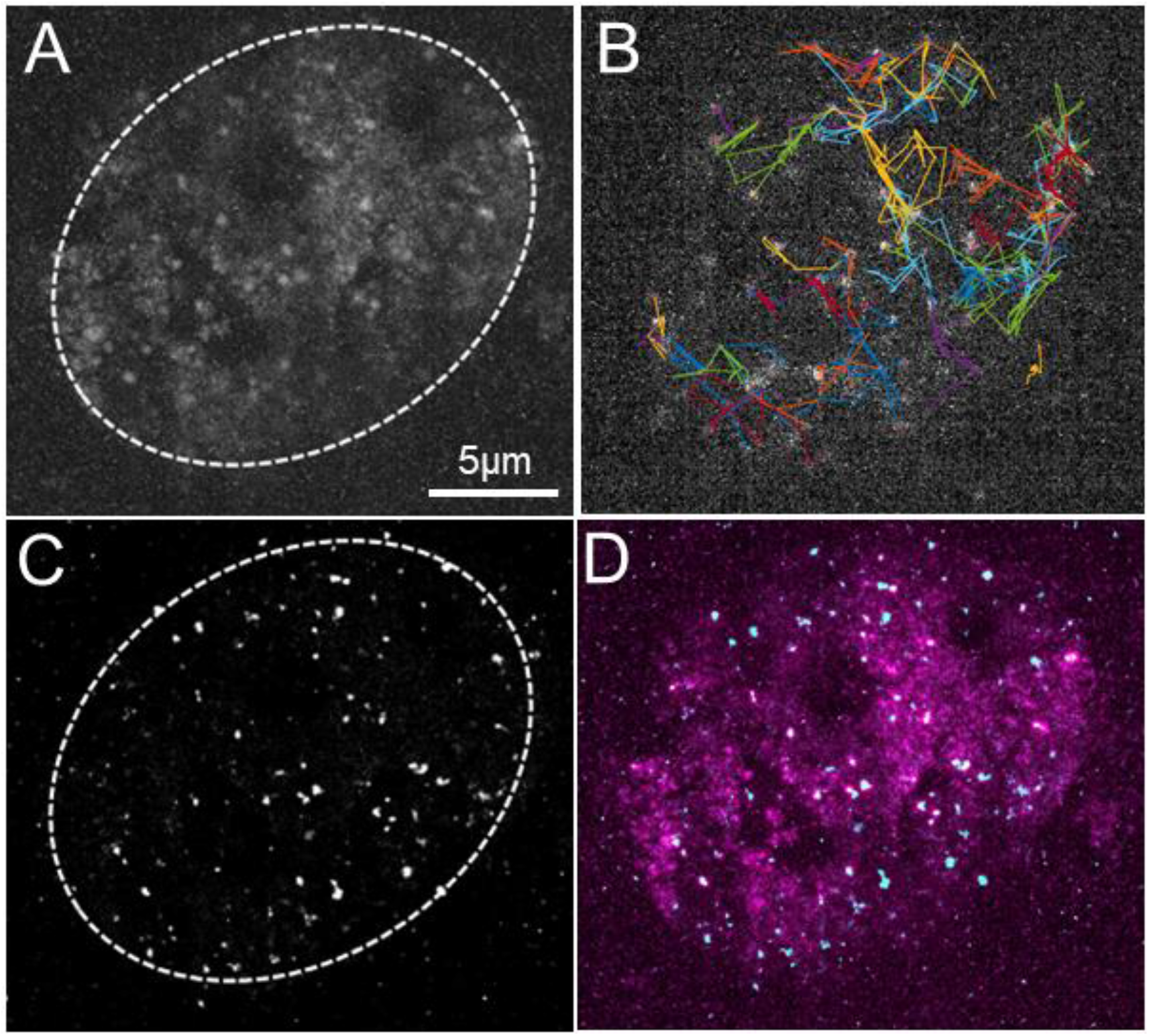
Spatio-temporal localization of RPB1 subpopulations. These images were acquired as a separate experiment within a 10s movie, after staining with JF549 at 1 nM. Panel A displays the mean intensity projection throughout the movie. Panel B illustrates the trajectories tracked at a given frame within the movie. Panel C represents the projection of all detected localizations using MTT in the whole film. Picture D merges the information from pictures A (in purple) and C (in cyan). The movie is also provided in supplementary data (Movies S3, S4 and S5), with raw data (S3), ongoing tracking superimposed (S4), or detections summed over time (S5).

Altogether, these data point out a very dynamic orchestration of RPB1 diffusion and activity.

## DISCUSSION

Quantifying the mobility of molecules in living cells is a very important issue for understanding the regulation of physiological mechanisms. The FCS and SPT techniques are highly suitable for such studies. However, because they strongly differ in acquisition methods, analysis, and experimental parameters, they are not easily comparable. In addition, the photo-physical properties of fluorophores (i.e. blinking, lifetime, etc.) affect the two techniques in different ways. For these two techniques to produce complementary information, it is necessary to properly characterize their windows of sensitivity and identify their limitations.

FCS is known to be particularly suited for the quantification of fast diffusion events but the classical ACF analysis often misses low-frequency events, whereas SPT allows the study of diffusion heterogeneities on a molecule-by-molecule basis, and potentially gives access to the monitoring of rare molecular events. One main limitation of SPT remains its low sensitivity to fast displacements because a large number of photons must be detected to allow accurate localization of single molecules, which requires integration times in the millisecond (ms) scale, while FCS is using sampling in the microsecond scale (μs). However, labeling with bright fluorophores presently allows obtaining good frame rates and track particles for tens of time points ^36^. The optical settings and configurations used in FCS or SPT are also quite different, with each instrument response functions (IRF) that must be considered to define their influence.

We, therefore, reasoned that a measurement-driven comparison between the two techniques was instrumental. For that purpose, we first characterized the influence of the setups on the diffusion measurements of calibrated beads. Whereas the use of simple models such as beads is widespread in FCS characterization, few works have dealt with this concept in SPT. We selected this solution, over *in silico* simulations of random behavior generation, because this approach displays the advantage of taking into account instrumental functions and real experimental noise. Thus, we could compare the performances of both techniques with relatively simple experimental settings. We show that SPT gives reliable results with high accuracy for diffusion coefficients below 4 μm^2^/s (Tt > 0.25 s/μm^3^), while FCS allows measurements from 1 to 15 μm^2^/s (Tts < 0.33 s/μm^3^).

To better control the measurement and analysis parameters of heterogeneous samples, we developed a benchmark based on a mixture of tracks from bead populations with different diffusion coefficients. Interestingly, our SPT analysis workflow, relying on the result distribution fitting by a Gaussian Mixture Model (GMM), discriminated quite well between the two initial subpopulations and obtained values very close to the expected ones (i.e., measured in the initial mono-populations). Moreover, the proportions of populations given by the analysis were also close to those used to make the mixtures, with very low variability. Taken together, we can consider that the proportions are reliable with a relative uncertainty as low as 10 %.

We then applied this validated workflow coupling SPT and FCS to the elucidation of the RNA polymerase II choreography in cell nuclei. Labelling of the RPB1 subunit of polymerase II (RNAP II) with fluorescent proteins, or Halo tags and adequate ligands, allowed us to measure RPB1 molecular dynamics. The analysis of RNAP II SPT trajectories with H-MSD showed that 20 % of RPB1 molecules appeared immobile for more than 60ms. Among RPB1 trajectories, we discriminated at least two main populations by SPT/H-MSD. The fastest population, representing approximately 20 % of RNAP II particles, displayed a mean transit-time (Tt) of 4.2 s/μm^3^ (D_60ms_ =0.5μm^2^/s), and an anomalous coefficient very close to 1 (mean ɑ=0.95), which reflects fairly well a free Brownian diffusion. This value is in the same order of magnitude as those found for the free diffusion of transcription factors ^23,37^, but it is an order of magnitude lower than the measurements of GFP multimer diffusion in the nucleus ^38^. This could indicate that free RPB1 and transcription factors have a slowed down Brownian diffusion in the nucleoplasm. The second mobile population that we identified by SPT/h-MSD had a mean transit-time (Tt) of 777 s/μm^3^ (D_60ms_ =0.02 μm^2^/s), and a mean anomalous coefficient ɑ=0.31, and represented 60 % of particles. This RNAP II subpopulation thus displayed a highly constrained subdiffusion and a very slow motion, even if it was distinct from bound immobile RNAP II.

We also established that most RPB1 diffusion measurements by FCS exceeded the speeds caught by SPT in our benchmark, even though such molecules probably contributed to the background noise of SPT acquisitions. RPB1 mean diffusion measured by FCS was characterized by Tt=62.10^−3^ s/μm3 (D_ɑ_=5.7 μm^2^/s^α^) and α= 0.61. These values are very close to those that we previously observed for CYCT1 ^39,40^, which suggests that these RPB1 mobility measurements may correspond to RNAP II molecules involved in the transcription pause release process. FCS, which is more sensitive to fast diffusions, hardly detected the subpopulations analyzed in SPT, with most molecules displaying transit-times 2 to 4 orders of magnitude higher than those identified by FCS, thus highlighting the complementarity of the two approaches.

The FCS data histogram distribution was imperfectly fit by Gaussian functions and displayed an array of relatively fast-diffusing molecules with a continuous range of diffusion parameters. We assume that within this range, there are no strict categories of RPB1 behaviors but rather transitions between states of interactions progressively evolving. Such behaviors may refer to the recent concepts that multivalent and dynamic interactions can drive phase separation ^41^ and that the function of large complexes could be maintained while molecules are constantly exchanged ^42^.

Interestingly, another spatiotemporal clustering analysis described foci predominantly occupied by a single RNAP II at the same time point ^43^, in agreement with data from immunogold labelling in live cells, concluding on the absence of significant RNAP II clustering ^45^. This was also recently confirmed with an investigation of RNA nanodomains that evidenced small RNAP II complexes mainly displaying only one RNAP II molecule ^46^. The absence of regions of high RNAP II concentration in our observations is thus completely consistent with these highlights. Still, we observed in our SPT movies subdomains oversampled by fast-moving of RNAPII. Quite remarkably, the population measured by FCS shown a motility with 3 to 6 magnitude order greater than which measured in other subpopulation (fig S4) and subdiffusion (α=0.61).To understand the mobility of RNAP II involved in this process we can think of the dynamics of a ball in a pinball machine, which presents very fast movements in a constrained space which increases the probability of encountering its targets. When these oversampling movements are observed with a sufficient integration time (30s-1 min) we obtain the image of a stronger concentration of fluorescence in these bumper-like domains. We hypothesize that this could be a mechanism for RNAP II recruitment and regulation of transcription. It is interesting to note that CYCT1, a cofactor of the transcriptional pause release, was also described to oversample the nuclear environment^3^. Moreover, we previously highlighted that CYCT1displays very similar diffusion properties, in subdomains whose size is in the same order of magnitude as that of TAD (Topologically Associated Domains) ^39,47^.. Taken together, our results thus reinforce the arguments in favor of the “RNAP II clustering” concept ^9^. The proper execution of RNAP II activity would rely on the increased probability of RPB1 to visit subdomains via an oversampling without necessarily increasing its concentration in a specific site (pinball-dumper effect). Interestingly, the study of Herpes Simplex Virus replication recently evidenced the possibility for RNAP II to visit in a repeated manner nearby DNA binding sites without clear boundaries between territories, which may be a common process ^50^. In that frame, highly dynamic interactions would ensure the transcription complex activity is maintained despite the exchange of molecules ^42^. Such a link between subdiffusion, topological frameworks and recruitment/regulation steps in transcription is emerging in the field ^48^ and starts to be described at the molecular level ^49^.

Altogether, our combined experiments suggest an RNAP II dynamic model with at least four kinetic subpopulations: a) chromatin-bound RNAP II, b) RNAP II slowly-diffusing in a Brownian manner, and two major subpopulations: c) subdiffusive slowly-diffusing RNAP II and d) fast-diffusing and oversampling RNAP II molecules. Moreover, our data also suggest the potential existence of other subpopulations. Future work will be aimed at elucidating whether we can identify and investigate these minority subpopulations to refine our knowledge of the transcription process.

In addition, FCS gave us access to molecular concentration in focal volume and indicated the presence in average of 2.6 +/−1.4 RPB1 molecules per focal volume, i.e. approximately 14 molecules per μm^3^. Considering the previous estimation of 80,200 RNAP II molecules per nucleus in U2OS cells, i.e. 37 molecules per μm^3^ ^43^, this would mean that FCS detected almost one-third of RPB1 molecules. In a first approach, we can extrapolate that the remaining two-thirds of RNAP II molecules, 23 out of 37 molecules per μm^3^, are observed by SPT. Within this category, 20% of SPT trajectories (i.e. 13 % of total RNAP II) were immobile (7-8 molecules/μm^3^). This estimate stands in agreement with the number of chromatin-bound RNAP II proposed by Cisse et al with another microscopy method ^9^. Besides, about 40% of RNAP II displayed a strong sub-diffusive behavior (mean anomalous coefficient ɑ=0.31). Taken together, these data point out a global view of a majority of RNAP II molecules that are strongly restricted, and potentially involved in regulatory steps. In SPT, only 20% of trajectories (i.e. 13 % of total RNAP II) could be considered free, according to their anomalous coefficient ɑ=0.95. This is again in agreement with the literature, including recent FRAP measurements concluding on a minority fraction of freely diffusing polymerases ^44^. X

At this stage, we hypothesize that the observed mobilities are associated with the mechanism of recruitment of RNAPII to transcription regulation. We also hypothesize that the subpopulations described, on the basis of their mobilities, are corresponding to transient states of RNAPII molecules and not to stable molecular states. We propose two models of RNAPII recruitment that can account for the observations and measurements of RPB1 dynamics. Both models encompass a free population slowly diffusing in an almost Brownian walk. The first model displays a single type of cluster and two steps corresponding to successive stages of regulation (fig 7A): in a first step, RNAP II is “trapped” in a subdomain with a strong oversampling of this space (pinball-bumper effect), and in a second step it presents a very constrained diffusion and a very slow speed of displacement leading to its immobilization on chromatin. We can assume that it is then the first step corresponding to the entry into the topologically active domains (TADs) which induces the oversampling and the first level of sub-diffusion, while the association with the cofactors causes the second step. We can also think of a second model where RNAP II is trapped in two types of clusters via two distinct co-existing mechanisms (fig 7 B). In some clusters, significant oversampling could correspond to “RNAP II trapping” in gene regions controlled by transcriptional pausing (similarity with CYCT1), whereas in other clusters “RNAP II trapping” would induce a strong and constrained diffusion that may correspond to transcriptional regulations without transcriptional pausing.

**Figure 7.**
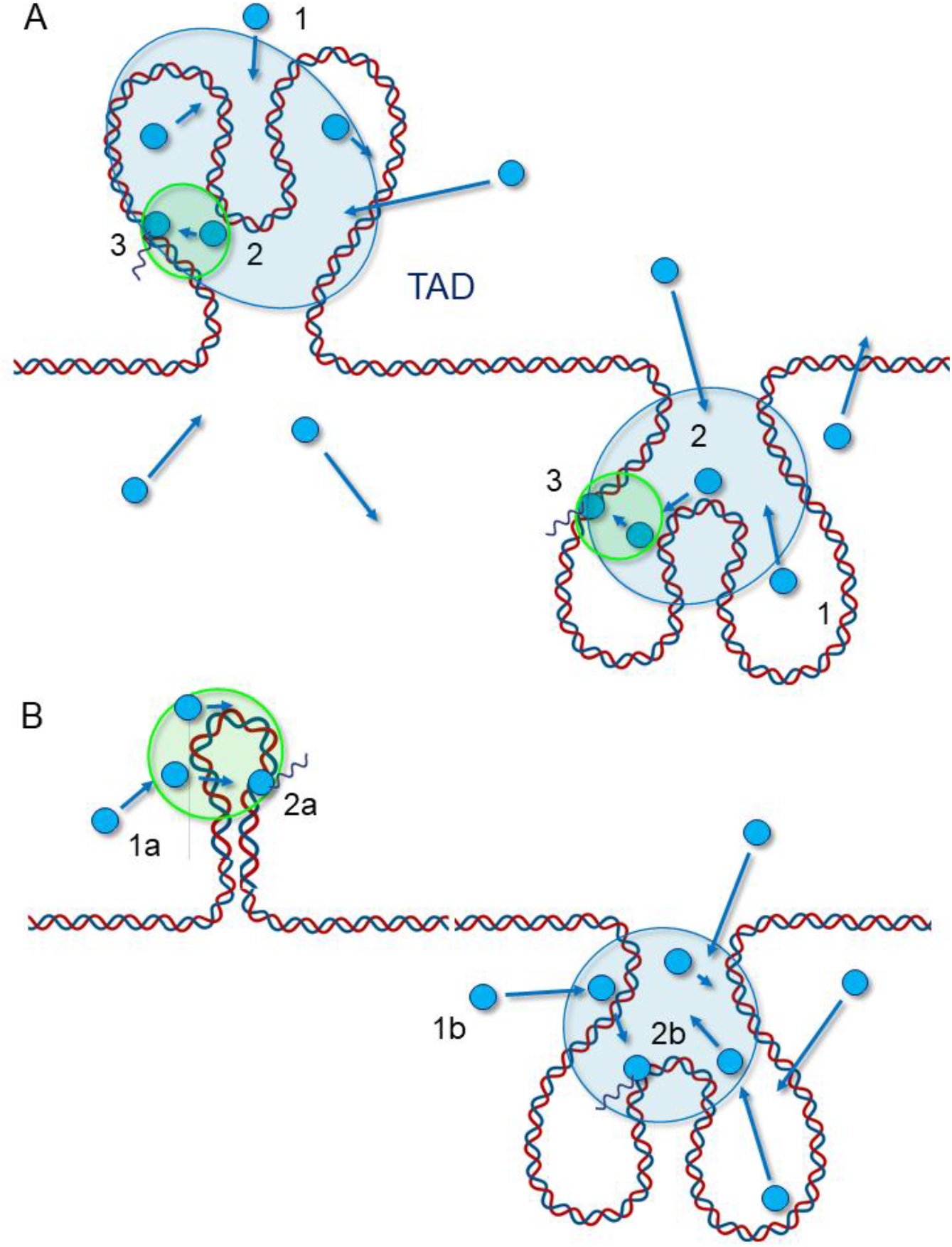
Models for RNAP II recruitment. This scheme displays the two hypotheses proposed to explain RNAP II mobilities and oversampling. The blue disks and arrows symbolize RNAP II molecules and their displacement vectors, respectively. A. Model with successive steps: 1) A free RNAP II (ɑ=0.95) enters and oversamples a large domain (blue), potentially a TAD, where it is confined (ɑ=0.61); 2) it is then trapped in a subdomain (green) of higher confinement (ɑ= 0.31), possibly through interactions with transcription factors, and binds to chromatin, 3) before being stabilized and starting transcription (fixed RNAP II). B. Model with distinct confinement domains (a and b) : 1a) and b) A free RNAPII (ɑ=0.95) can enter two types of confined domains, leading to fast oversampling or not; 2a) and b) RNAPII is then recruited and binds to chromatin (fixed RNAPII). This second model helps to account for the existence of different mechanisms of RNAPII recruitment and in particular the existence of the proximal transcriptional pause for only half of genes.

This study paves the way for future work that will be aimed at investigating which of these two hypotheses correspond to the reality of RNAP II recruitment to active genes, and at identifying the molecular mechanisms responsible for RNAP II subdiffusion. In that frame, the coupling of FCS and SPT acquisitions on the same instrument would represent an interesting instrumental development to simultaneously acquire molecules by both methods in a spatio-temporal manner, and we are moving forward in that direction.

## METHODS

### Reagents and sample preparations

#### Standard samples preparation

Water/Glycerol solutions were prepared considering weight percentage with the use of a precision balance. Solutions were vortexed for 30 sec before being stored at 37°C. A few minutes before acquisitions, to avoid agglomerations of beads, we diluted our fluorescent beads stocks in two distinct manners:

- 100 nm diameter beads (Molecular Probes, Fluorescent microspheres (505/515), 2 % solid) diluted 10^4^ in solutions for SPT acquisition
- 40 nm diameter beads (Molecular Probes, Fluorescent microspheres (505/515), 2 % solid) diluted 10^2^ in solutions for FCS acquisition

To compare the SPT and FCS workflows, we applied a similar strategy. However, since the experimental focal volume in FCS was in the order of 200 nm diameter in the x-y axis, the use of 100nm-diameter beads resulted in huge bursts of fluorescence during signal acquisition, which were misinterpreted during the Autocorrelation Function (ACF) process. We therefore chose smaller fluorescent beads (40nm-diameter) for FCS experiments, which allowed us to measure more reliably fluorescence fluctuations and infer their diffusion coefficients.

Bead solutions were finally deposited in 35-mm glass-bottom dishes (Ibidi, Clinisciences, France) and kept at 37° in a thermostatic chamber during acquisitions.

The theoretical values of beads’ diffusion were calculated according to the Stokes-Einstein formula:

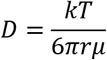

With *k* Boltzmann constant, T temperature in Kelvin, r the hydrodynamical radius of the bead and μ the viscosity.

Viscosity values of different (w:g) solutions were calculated following Takamura et al. ^26^ and are indexed in the following table.

| % glycerol | Calculated D value ($\mu\text{m}^2/\text{s}$) | Calculated D value ($\mu\text{m}^2/\text{s}$) |
| --- | --- | --- |
|  | for 40 nm beads | for 100 nm beads |
| 0 | 15.1 | 5,98 |
| 10 | 12.1 | 4,78 |
| 20 | 9.05 | 3,58 |
| 50 | 3.24 | 1,25 |
| 80 | 0.4 | 0,19 |

### Biological resources

#### Cell line culture

Human U2OS osteosarcoma cells (HTB-96 accession number from the American Type Culture Collection, Manassas, VA) were tested for mycoplasma contamination before establishing the RPB1 cell lines. Cells were cultured in DMEM medium (Gibco Laboratories, Gaithersburg, MD) supplemented with 10% FCS (V/V) and penicillin/streptomycin (100 mg/mL). Cells were grown in a humidified incubator at 37°C / 5 % CO2. Cells were passaged every 2–4 days before reaching confluency.

We generated plasmids encoding for human RPB1 (N792D) α-amanitin-resistant mutated fused to an N-terminal Halo tag, by molecular cloning.

We stably transfected U2OS cells with this plasmid using Fugene 6 following the manufacturer’s instruction (Promega). α-Amanitin (SIGMA #A2263) was used to select transfected cells, at a concentration of 2 μg/mL and was used thereafter in permanence in the culture of the cells at a concentration of 1μg/mL to avoid endogenous RPB1 re-expression as described in (Boehning et al. 2018). For live-cell imaging, cells were plated on 35-mm glass-bottom dishes (Ibidi, CliniSciences, Nanterre, France), filled with L-15 medium without Phenol Red (Life Technologies, Carlsbad, CA), and incubated at 37°C in a microscope thermostatic chamber (OKO, Italia) for imaging.

#### Labeling protocol

We labeled cells just before imaging. Following HaloTag technology protocol (Promega), cells were incubated for 30 min in DMEM 10%FCS supplemented with 0.1 nM JF 549 for SPT. Then, cells were rinsed 3 times with DMEM + 10% SVF and were re-incubated for 30 min. Finally, the cell medium was replaced by L-15 medium (Life Technologies, CA, USA) before imaging, to avoid DMEM auto-fluorescence.

#### SPT set up

SPT measurements were performed with an inverted microscope (Ti-Eclipse, Nikon Instrument, Japan) adapted with an x100 NA=1,49 TIRF water immersion objective and an appropriate dichroic 405/488/561/635 nm (ref Di03-R405/488/561/635 SEMROCK, USA). The excitation laser (488nm Argon LASER, Melles-Griot, USA and 561nm, OXXIUS, France) was injected into the TIRF Nikon module into a mono-mode fiber and focused on the backplane of the objective. Illumination was performed in HILO mode ^29^ to improve contrast in images. Perfect Focus System (PFS, Nikon) was set to avoid drift on Z-axis (defocusing) on the objective, relative to the coverslip. Experiments were acquired under continuous illumination (488 nm 20 kW/cm^2^ on the sample, or 561 nm 20 kW/cm^2^ on the sample). Emitted fluorescence was harvested on an sCMOS camera (PCO EDGE 4.2, PCO, Germany) with a pixel size of 6,5 μm. The acquisition format was 100×100 px, and the acquisition rate was set at 100 Hz (10 ms per frame). All devices were driven by NIS software (Nikon Instruments, Japan). A thermostatic chamber (‘on-stage-incubator’, OKOLAB, Italy) was used to keep samples at 37 Celsius degrees.

#### SPT data analysis

Tracks were reconstructed (detection and tracking) with MTT algorithm tracking software ^30^ running on MATLAB (MathWorks), which is well adapted for a high concentration of particles (up to 10 for a 100×100px field of view). For each condition, we had at least 1000 trajectories lasting more than 10 detections (smaller trajectories were not included in the analysis). Tracks were then analyzed with custom code kindly provided by Laura Caccianini (Coppey/Hajj lab, Institut Curie, France). H-MSD (histogram Mean Square Displacement) technique is based on the Time-Average MSD calculation *TA* − *MSD*(*τ*) = ⟨*r*^2^(*τ*)⟩*T* = ⟨[*x*(*t* + *τ*) − *x*(*t*)]^2^ + [*y*(*t* + *τ*) − *y*(*t*)]^2^⟩_*T*_ ∝ 4*Dτ*^*α*^ where Δ*t*, with n the number of frames considered for the sliding average (0<n<*L_traj_*, *L_traj_*the length of the trajectory), Δ*t* the lag time between two frames (10 ms), D the diffusion coefficient (μm^2^/s), and α is the anomaly coefficient.

With H-MSD each track was treated individually as described in ^32,51^. The TA-MSD curves were fitted with a linear fit until *τ* = 60 *ms* to obtain the instantaneous diffusion coefficient (*D*_60*ms*_) following the previous works of Michalet et al. describing a procedure to analyze the individual MSD curves ^31^. In accordance with this work, the first point MSD (1Δ*t*) is not considered in the fit because it is the same order of magnitude as the static localization error (50 nm, calculated on fixed RPB1-Halo in U2OS cells.

Anomaly coefficient alpha (α) was calculated by fitting the log-log representation of the TA-MSD with a power-law function TA-MSD(t)∝ 4*t*^*α*^. For individual tracks, it is common that the second half of the MSD curve is noisy because of the low statistics for each MSD(nΔ*t*) value. We decided to consider the first third of the curve for the linear fit because more than the second half of the MSD curve is noisy and cannot be interpreted. The slope of the regression line gave alpha (α). In addition, the diffusion coefficient could also be computed from the intercept of the slope, D_α_, and was compared to D_60ms_ in the results.

A Gaussian mixture model (GMM) was applied to fit the data. GMM was used as a clustering algorithm where the number k of populations is chosen from an analysis of the Bayesian Information Criterion and the Akaike Information Criterion. This population number was chosen as small as possible, following those criteria. By evaluating the number of discernible populations with a statistical criterion we tried to minimize any bias that can occur when using such a supervised machine learning algorithm and create a robust and reproducible analysis pipeline.

#### Mix diffusion data models

For the two sub-populations studies, we mixed batches of trajectories coming from different conditions. The different conditions and the different fractions for each mix are indexed in the following tables.

**Combined SPT and FCS methods highlight a mechanism of RNAP II oversampling in cell nuclei**
| Condition A |  | Total number<br>of trajectories | Nb of D1<br>trajectories | % D1 | Nb of D2<br>trajectories | % D2 |
| --- | --- | --- | --- | --- | --- | --- |
| D1=4 $\mu\text{m}^2/\text{s}$<br>(w :g =100 :0) | m1 | 2013 | 845 | 42 | 1168 | 58 |
| & D2=0.1 $\mu\text{m}^2/\text{s}$<br>(w :g=20 :80) | m2 | 3118 | 2339 | 75 | 780 | 25 |
|  | m3 | 3369 | 842 | 25 | 2527 | 75 |

| Condition B |  | Total number<br>of trajectories | Nb of D1<br>trajectories | % D1 | Nb of D2<br>trajectories | % D2 |
| --- | --- | --- | --- | --- | --- | --- |
| D1=0.9 $\mu\text{m}^2/\text{s}$<br>(w :g =50 :50) & | m1 | 2027 | 932 | 46 | 1095 | 54 |
| D2=0.1 $\mu\text{m}^2/\text{s}$<br>(w :g=20 :80) | m2 | 3207 | 1860 | 58 | 1347 | 42 |
|  | m3 | 3633 | 908 | 25 | 2725 | 75 |

| Condition C |  | Total number<br>of trajectories | Nb of D1<br>trajectories | % D1 | Nb of D2<br>trajectories | % D2 |
| --- | --- | --- | --- | --- | --- | --- |
| D1=4.0 $\mu\text{m}^2/\text{s}$<br>(w :g =100 :0) & | m1 | 4470 | 894 | 20 | 3576 | 80 |
| D2=0.9 $\mu\text{m}^2/\text{s}$<br>(w :g=50 :50) | m2 | 4791 | 2443 | 51 | 2348 | 49 |
|  | m3 | 5119 | 3993 | 78 | 1126 | 22 |

| Condition D |  | Total number<br>of trajectories | Nb of D1<br>trajectories | % D1 | Nb of D2<br>trajectories | % D2 |
| --- | --- | --- | --- | --- | --- | --- |
| D1=4.0 $\mu\text{m}^2/\text{s}$<br>(w :g =100 :0) & | m1 | 5293 | 688 | 13 | 4605 | 87 |
| D2=3.8 $\mu\text{m}^2/\text{s}$<br>(w :g=90 :10) | m2 | 5310 | 2443 | 46 | 2867 | 54 |
|  | m3 | 5398 | 4049 | 75 | 1350 | 25 |

#### FCS set up

FCS measurements were performed on a Nikon A1 confocal microscope (Nikon Instrument, Tokyo, Japan) with a 40x objective NA=1,25 (Nikon Instrument, Tokyo, Japan). Excitation Argon LASER 488 nm (Coherent, USA) was used to scan samples in confocal illumination. To spectrally select the excitation from the emitted light, we used a bandpass dichroic mirror (405/488 nm). Fluorescence photons were collected on a photodiode and images were reconstructed with NIS software (Nikon Instruments, Tokyo, Japan).

To perform FCS, the CLSM was adapted with a TCSPC module (PicoQuant, Germany), and a pulsed laser at 488 nm (LDH-D-C 488, PicoQuant), a TSCPC unit, and a single molecule counting detector (SPCM CD 3516H, Excelitas Technologies Corp., USA). To drive this photon counting system, we used SymphoTime software (PicoQuant, Germany) in FCS mode. A high pass filter (BLP-488R, SEMROCK, USA) was set before the detector.

#### FCS analysis

FCS setup was calibrated with ATTO 488 diluted at 10 nM in Milli-Q water to define the focal volume by best practice. In the water at 36°, the temperature of our incubator, the Atto 488 diffusion coefficient is 536 μm^2^/s (source: PicoQuant Application Note Kapusta and GmbH, n.d.).

The number N of molecules in this observation volume V was inferred from the autocorrelation function with

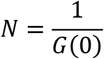

And the observation volume determined with:

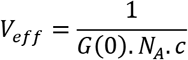

N_A_ is the Avogadro number and c is the concentration of the calibration solution.

Once calibration was done, we had a good estimate of the focal volume of the microscope and could proceed to the analysis of fluorescence fluctuations. Measurements displaying a linear regression of the fluorescent signal characteristic of photobleaching were excluded. For data analysis, we use a home-written MATLAB code named EASY FCS ^40^ which can fit the autocorrelation function (ACF).

In the case of normal diffusion the ACFs were adjusted with a Gaussian function (assuming the beam as a Gaussian profile) :

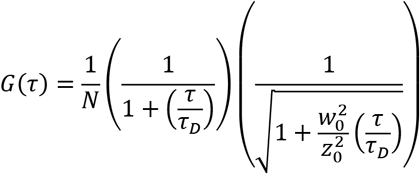

Where *N* is the number of molecules and *τ_D_* is the characteristic residence time.

For anomalous diffusion, the mean square displacement is no longer linearly dependent on time, but it evolves proportionally with time to the power of α. In this case, the autocorrelation function becomes:

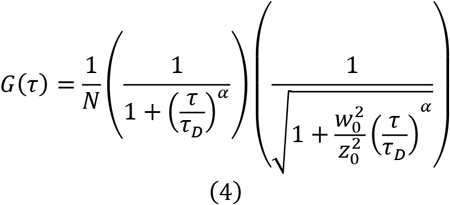

The fitting of the ACFs was done with the Trust-Region-Reflective Least Squares Algorithm. To evaluate the goodness of the fit, we minimized the sum of the squared error.

Since the diffusion coefficient (D) obtained by anomalous model fit is given with time dimensions dependent on exponent α, it is recommended to give the results in transit time ^23,24,39^ to compare data.

#### Standardization of analysis metrics between FCS and SPT

FCS and SPT data are obtained by analytical methods from microscopic approaches with different metrics. In order to compare them and use them together to analyze molecular mobility, it is useful to express them with the same metric and unit. The adjustment of the results by an anomalous model in both techniques also induced a difficulty, because when the anomalous coefficient changed, the time unit was simultaneously modified. To overcome this difficulty, it is usual in FCS to express the ACF results as residence times in the focal volume. Here, we propose to extend this metric to SPT data by proposing a transit time (Tt) within a reference volume (s/μm^3^) which could then be a common metric to both FCS and SPT techniques. In a first approach, we make the hypothesis that the displacements measured by SPT are isotropic in the volume at the scale of the confocal resolution (femtoliter), to pass from a surface measurement to the exploration of the corresponding volume. This allows us to define a normalized transit time from each measurement.

## ACKNOWLEDGEMENTS

We thank all members of our team and GDR Imabio-CNRS, particularly Dr Hugues Berry, Dr Ignacio Izeddin and Dr Thorsten Wohland for their help and fruitful discussion/comments. We are much grateful to Dr Raoul Torero-Ibad for his great work and a precious help in providing us with a functional lab environment, and for his enlightened opinion on this study. We thank Christian Hubert (Errol company) for his valuable input in instrumental advances. We are much grateful to Dr Jonathan Grimm and Dr Luke Lavis (Janelia Farm, VA, USA) for kindly providing us with JF dyes to perform our experiments.

## AUTHOR CONTRIBUTIONS

MF, PL, GB, AF, and LH designed research. MF, PL, DC, and FA performed experiments. MF, PL, AL, GB, AF, and LH analyzed and interpreted the data. MF, PL, AF, and LH wrote the manuscript.

## DATA AVAILABILITY STATEMENT

Data and material are available on request to

## SUPPLEMENTARY DATA

Supplementary Movies are available at:

https://nextcloud.univ-lille.fr/index.php/s/aqXXqPsMj93DESq
https://nextcloud.univ-lille.fr/index.php/s/Da6m2W9oGPexfcY
https://nextcloud.univ-lille.fr/index.php/s/KjXnoYcG26AnFXL
https://nextcloud.univ-lille.fr/index.php/s/HQW5rXBSg78JHMw
https://nextcloud.univ-lille.fr/index.php/s/eFTsxyXaPnApE9C

## SUPPLEMENTAL MATERIAL

**Supplemental Figure 1.**
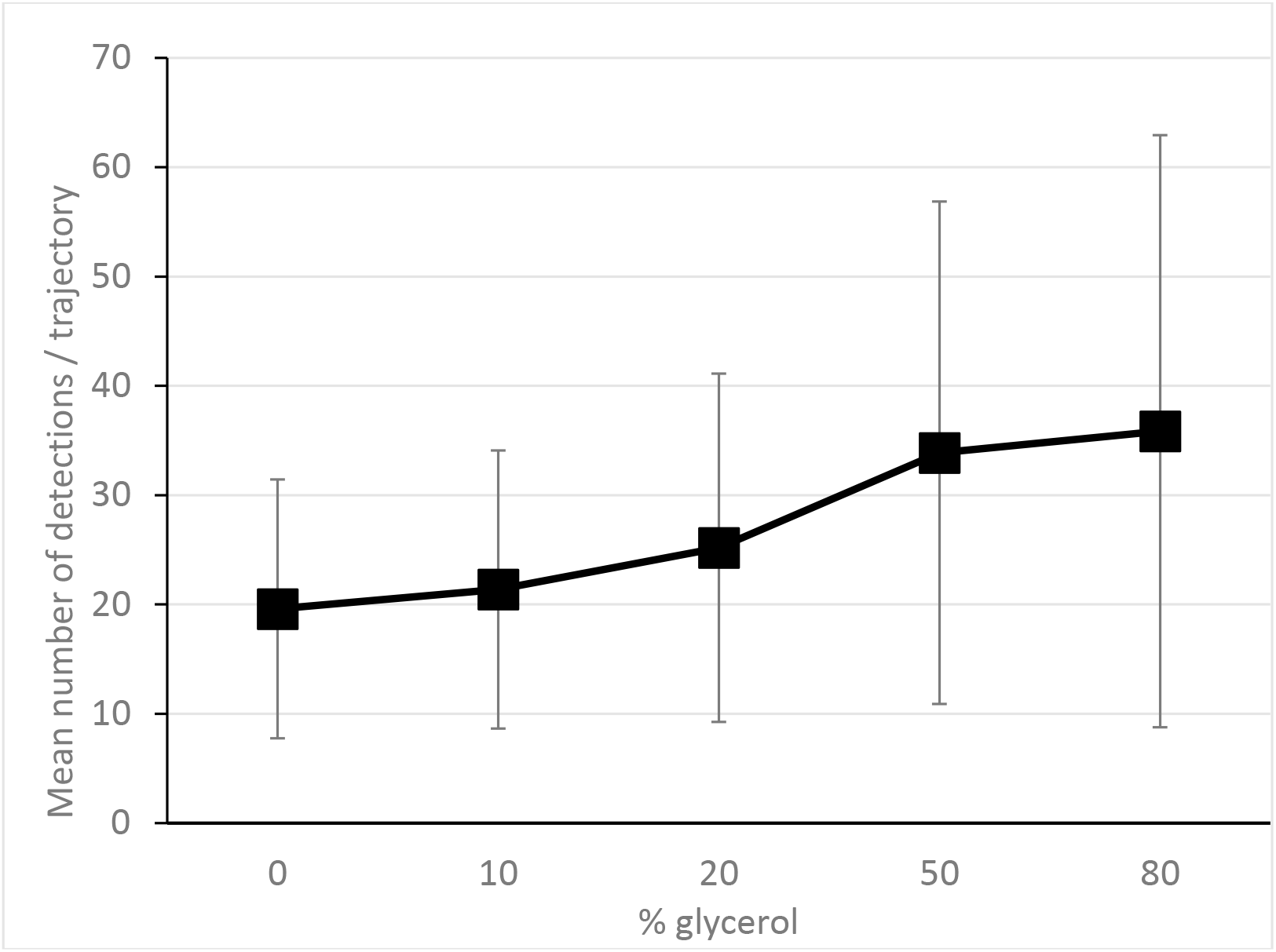
Mean number +/− standard deviation of detections obtained with Single Particle Tracking of fluorescent beads diffusing in aqueous solutions with different glycerol proportions.

**Supplemental Table 1.** Comparison of experimental vs theoretical values of anomaly coefficients (A) and diffusion coefficients (B) computed from the ACF of FCS measurements performed with fluorescent beads diffusing in aqueous solutions with different glycerol proportions.

| Anomaly coefficient | Experimental values |  | Theory |
| --- | --- | --- | --- |
| % glycerol | mean $\alpha$ | SD | $\alpha$ |
| 0 | 0.96 | 0.01 | 1 |
| 10 | 0.94 | 0.02 | 1 |
| 20 | 0.97 | 0.02 | 1 |
| 50 | 1.02 | 0.02 | 1 |
| 80 | 0.7 | 0.02 | 1 |

| Diffusion coefficient | Experimental values |  | Theory |
| --- | --- | --- | --- |
| % glycerol | mean $D_{\alpha}$ ($\mu\text{m}^2/\text{s}$) | SD | D ($\mu\text{m}^2/\text{s}$) |
| 0 | 15.5 | 1.7 | 15.1 |
| 10 | 12.1 | 1.8 | 12.1 |
| 20 | 9.2 | 1.5 | 9.0 |
| 50 | 4.5 | 0.8 | 3.2 |
| 80 | 1.1 | 0.2 | 0.4 |

**Supplemental Figure 2.**
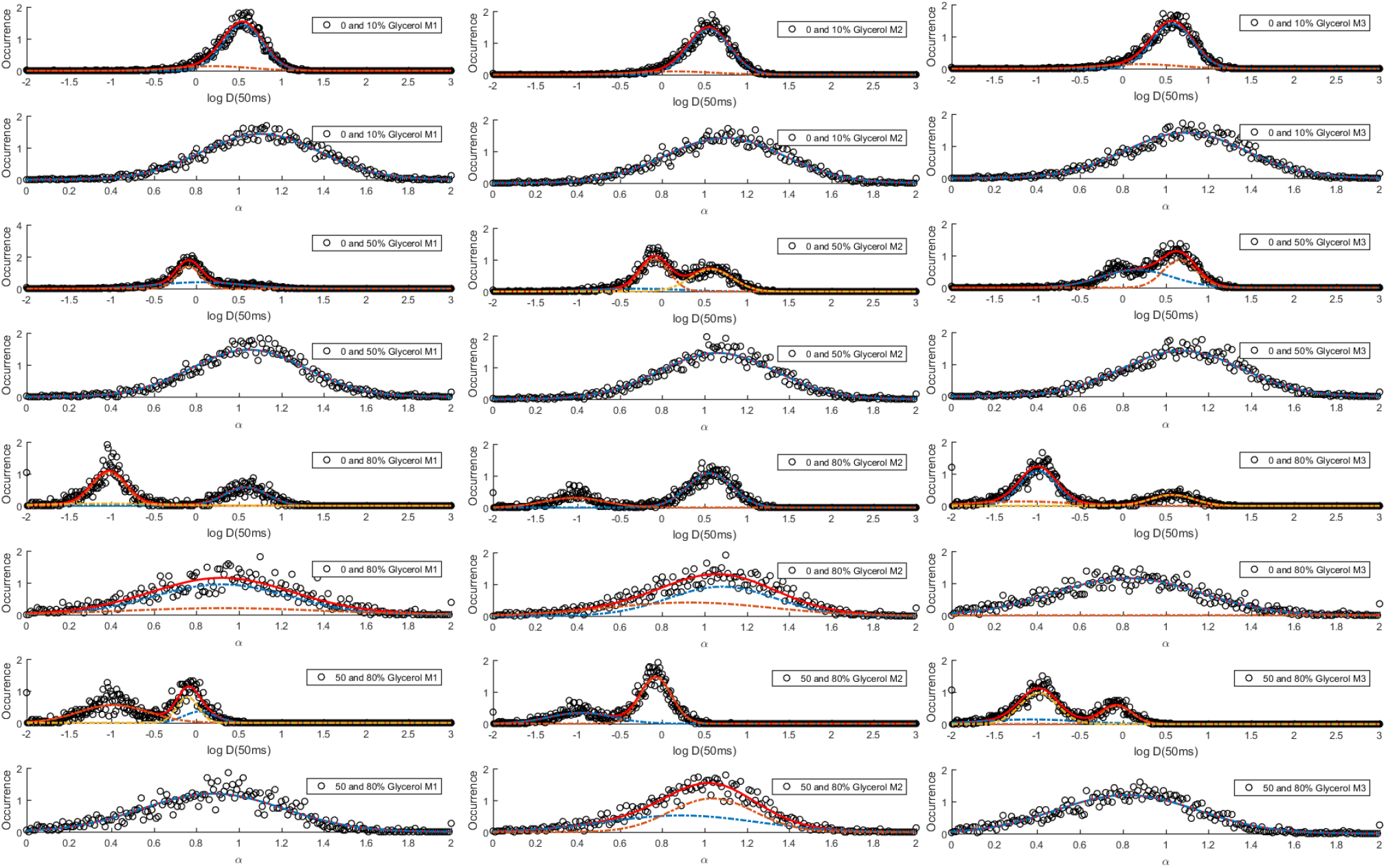
Representation of subpopulations identified through the GMM (Gaussian Mixture Model) after mixing batches of trajectories obtained in different conditions, as detailed within the graphics legend.

**Supplemental Table 2.**
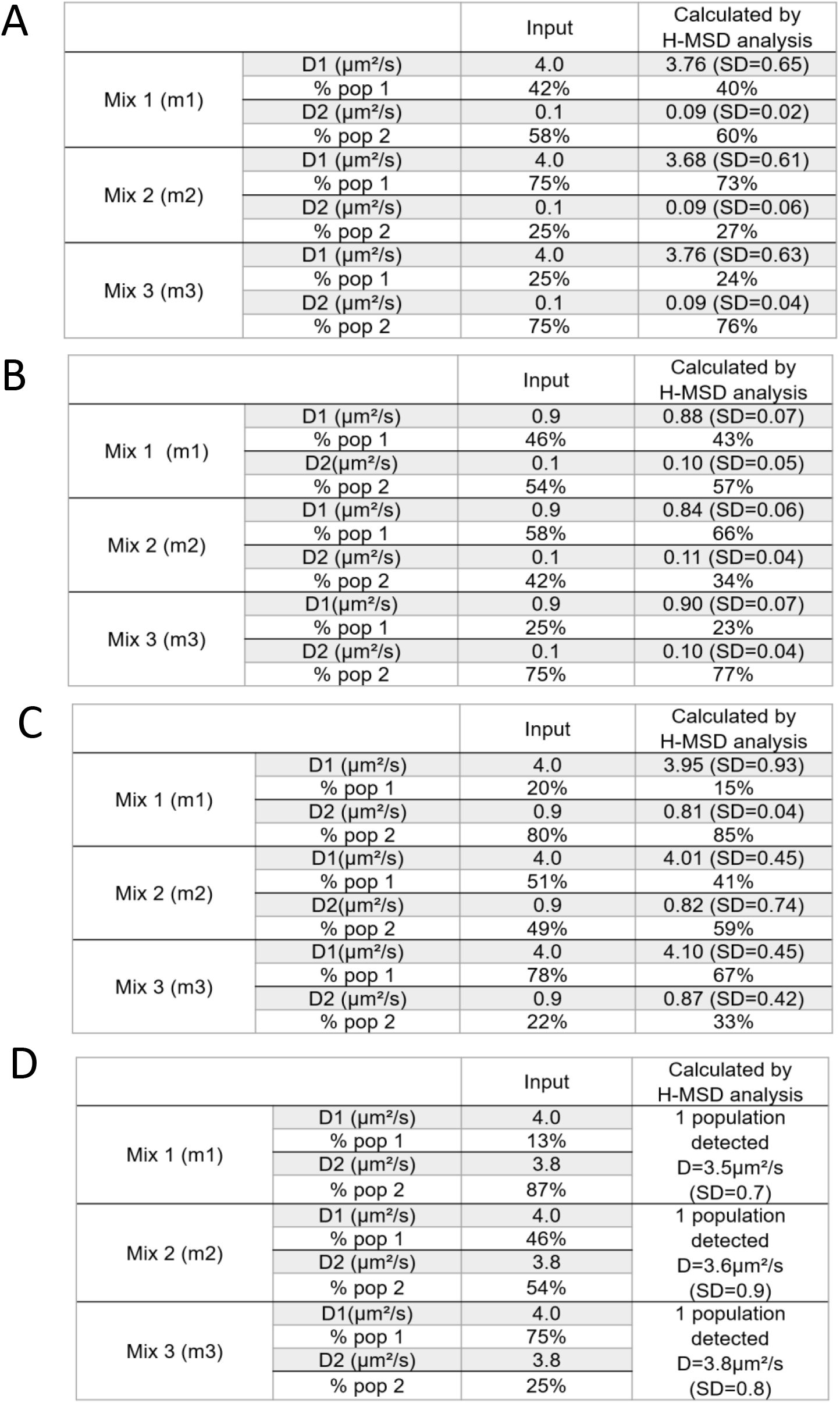
Comparison of input values vs h-MSD computed values after mixing batches of trajectories obtained in different conditions.

|  |  | Input | Calculated by<br>H-MSD analysis |
| --- | --- | --- | --- |
| Mix 1 (m1) | D1 ($\mu\text{m}^2/\text{s}$) | 4.0 | 3.76 (SD=0.65) |
|  | % pop 1 | 42% | 40% |
| | D2 ($\mu\text{m}^2/\text{s}$) | 0.1 | 0.09 (SD=0.02) |
|  | % pop 2 | 58% | 60% |
| Mix 2 (m2) | D1 ($\mu\text{m}^2/\text{s}$) | 4.0 | 3.68 (SD=0.61) |
|  | % pop 1 | 75% | 73% |
| | D2 ($\mu\text{m}^2/\text{s}$) | 0.1 | 0.09 (SD=0.06) |
|  | % pop 2 | 25% | 27% |
| Mix 3 (m3) | D1 ($\mu\text{m}^2/\text{s}$) | 4.0 | 3.76 (SD=0.63) |
|  | % pop 1 | 25% | 24% |
| | D2 ($\mu\text{m}^2/\text{s}$) | 0.1 | 0.09 (SD=0.04) |
|  | % pop 2 | 75% | 76% |

**B**
|  |  | Input | Calculated by<br>H-MSD analysis |
| --- | --- | --- | --- |
| Mix 1 (m1) | D1 ($\mu\text{m}^2/\text{s}$) | 0.9 | 0.88 (SD=0.07) |
|  | % pop 1 | 46% | 43% |
| | D2 ($\mu\text{m}^2/\text{s}$) | 0.1 | 0.10 (SD=0.05) |
|  | % pop 2 | 54% | 57% |
| Mix 2 (m2) | D1 ($\mu\text{m}^2/\text{s}$) | 0.9 | 0.84 (SD=0.06) |
|  | % pop 1 | 58% | 66% |
| | D2 ($\mu\text{m}^2/\text{s}$) | 0.1 | 0.11 (SD=0.04) |
|  | % pop 2 | 42% | 34% |
| Mix 3 (m3) | D1 ($\mu\text{m}^2/\text{s}$) | 0.9 | 0.90 (SD=0.07) |
|  | % pop 1 | 25% | 23% |
| | D2 ($\mu\text{m}^2/\text{s}$) | 0.1 | 0.10 (SD=0.04) |
|  | % pop 2 | 75% | 77% |

**C**
|  |  | Input | Calculated by<br>H-MSD analysis |
| --- | --- | --- | --- |
| Mix 1 (m1) | D1 ($\mu\text{m}^2/\text{s}$) | 4.0 | 3.95 (SD=0.93) |
|  | % pop 1 | 20% | 15% |
| | D2 ($\mu\text{m}^2/\text{s}$) | 0.9 | 0.81 (SD=0.04) |
|  | % pop 2 | 80% | 85% |
| Mix 2 (m2) | D1 ($\mu\text{m}^2/\text{s}$) | 4.0 | 4.01 (SD=0.45) |
|  | % pop 1 | 51% | 41% |
| | D2 ($\mu\text{m}^2/\text{s}$) | 0.9 | 0.82 (SD=0.74) |
|  | % pop 2 | 49% | 59% |
| Mix 3 (m3) | D1 ($\mu\text{m}^2/\text{s}$) | 4.0 | 4.10 (SD=0.45) |
|  | % pop 1 | 78% | 67% |
| | D2 ($\mu\text{m}^2/\text{s}$) | 0.9 | 0.87 (SD=0.42) |
|  | % pop 2 | 22% | 33% |

|  |  | Input | Calculated by<br>H-MSD analysis |
| --- | --- | --- | --- |
| Mix 1 (m1) | D1 ($\mu\text{m}^2/\text{s}$) | 4.0 | 1 population<br>detected<br>D=3.5$\mu\text{m}^2/\text{s}$<br>(SD=0.7) |
|  | % pop 1 | 13% |  |
| | D2 ($\mu\text{m}^2/\text{s}$) | 3.8 | |
|  | % pop 2 | 87% |  |
| Mix 2 (m2) | D1 ($\mu\text{m}^2/\text{s}$) | 4.0 | 1 population<br>detected<br>D=3.6$\mu\text{m}^2/\text{s}$<br>(SD=0.9) |
|  | % pop 1 | 46% |  |
| | D2 ($\mu\text{m}^2/\text{s}$) | 3.8 | |
|  | % pop 2 | 54% |  |
| Mix 3 (m3) | D1 ($\mu\text{m}^2/\text{s}$) | 4.0 | 1 population<br>detected<br>D=3.8$\mu\text{m}^2/\text{s}$<br>(SD=0.8) |
|  | % pop 1 | 75% |  |
| | D2 ($\mu\text{m}^2/\text{s}$) | 3.8 | |
|  | % pop 2 | 25% |  |

**Supplemental Figure 3.**
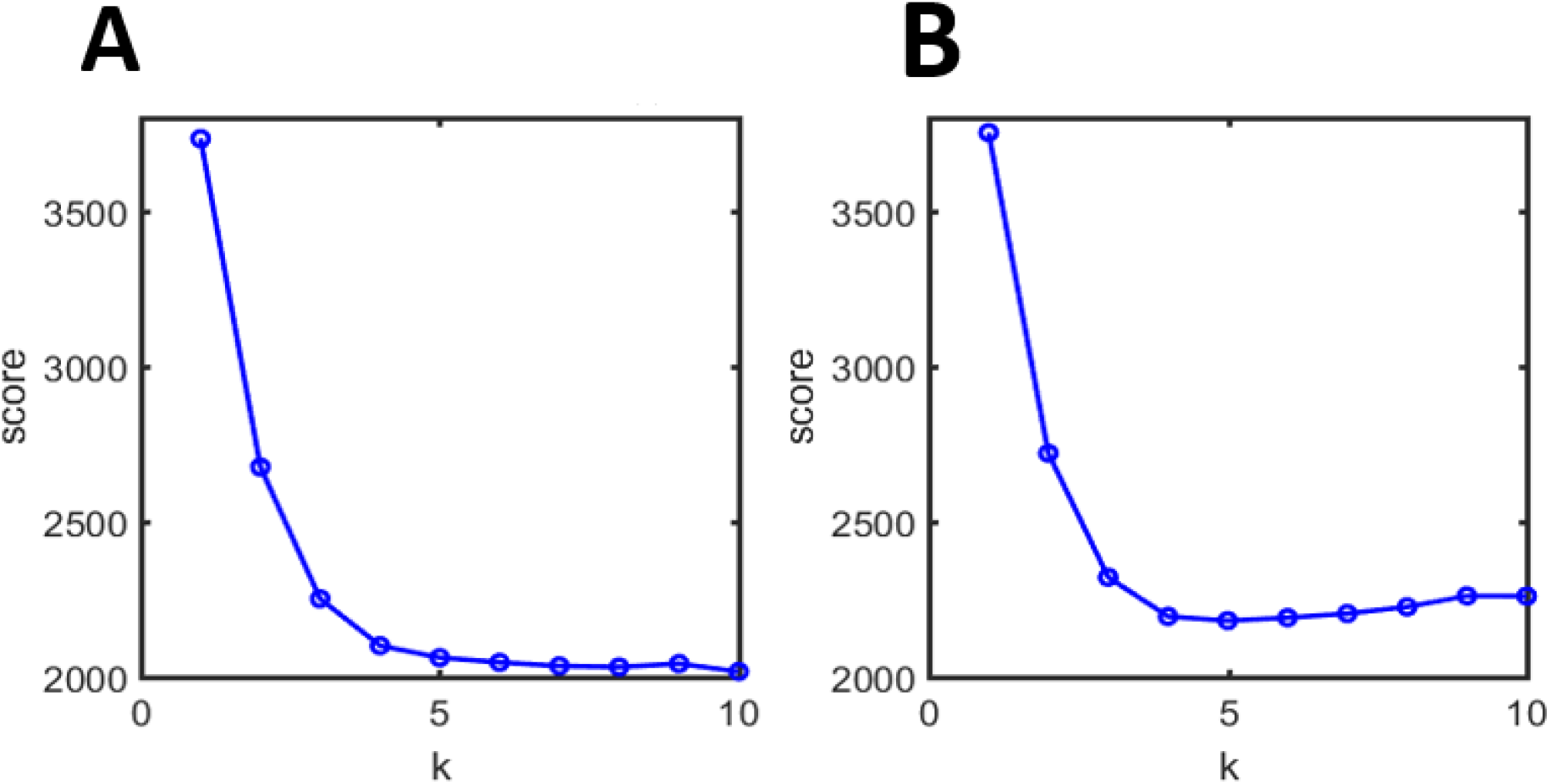
AIC (Akaike Information Criteria) (panel A) and BIC (Bayesian Information Criteria) (panel B) scores corresponding to the n-population analysis of RPB1 SPT trajectories

**Supplemental Figure 4.**
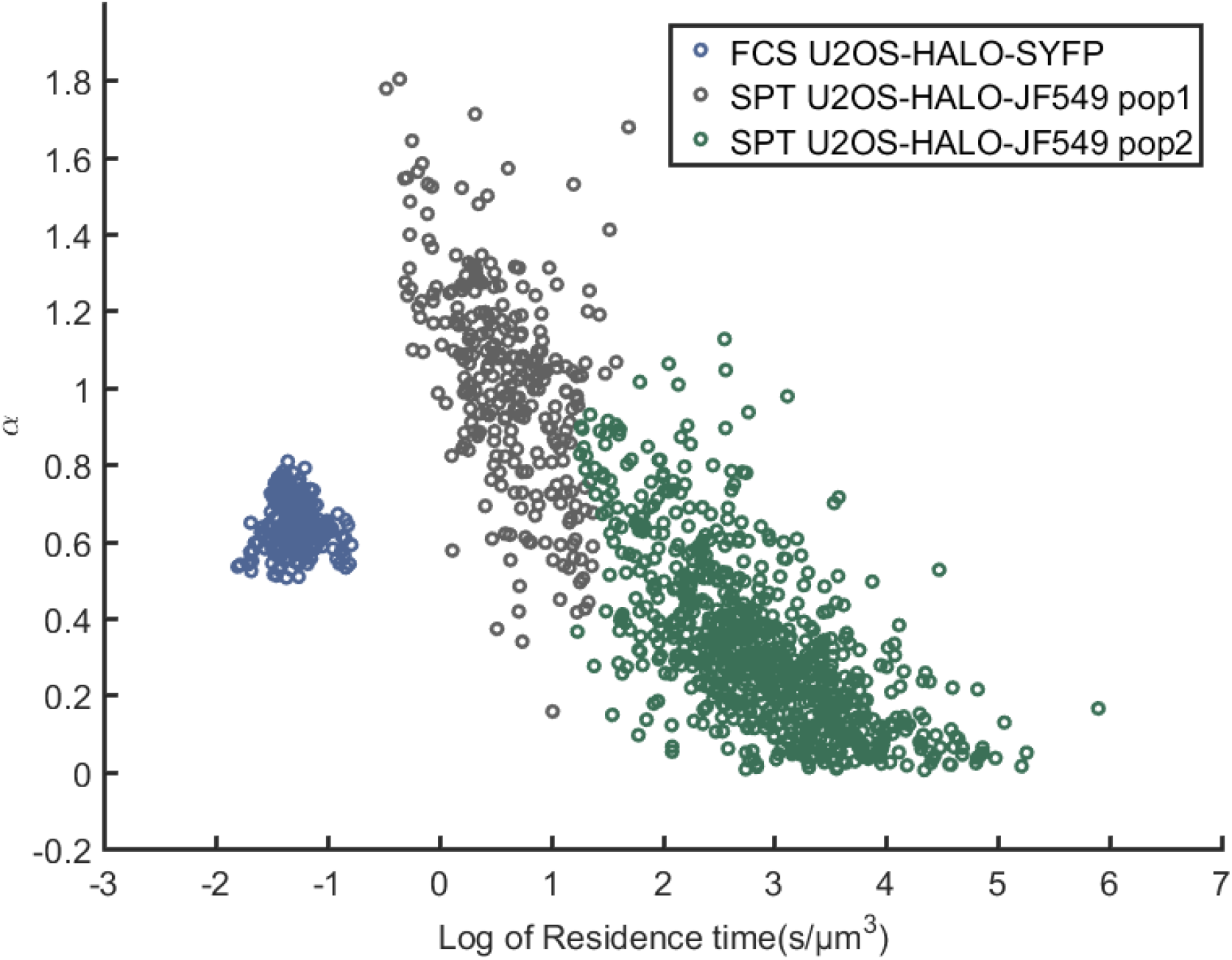
Diffusion parameters for RPB1 measured by FCS (blue dots) and SPT (green dots and gray dots, respectively corresponding to the main and minority subpopulations detected) are gathered in this scatter plot.

